# Can the Human Plastisphere Account for the Rise of Type 1 Diabetes: A Computational Molecular Mimicry Study

**DOI:** 10.64898/2026.11.02.763715

**Authors:** Julio C. Facelli, McKay Jones, Sejal Mistry, Ram Gouripeddi

## Abstract

An increased incidence of childhood type 1 diabetes mellitus (T1DM) has been widely documented, but its pathophysiological origins are poorly understood. The post-World War II expansion of the plastics industry in the United States and elsewhere coincides with a marked increase in T1DM incidence; however, no pathophysiological mechanism has been proposed that could justify such a temporal correlation. Here, we leverage recent advances in artificial intelligence and protein structure prediction and binding to show that molecular mimicry is a plausible pathophysiological mechanism for plastic fragments to induce onset of T1DM for individuals with certain HLA haplotypes.

## Introduction

An increased incidence of childhood type 1 diabetes mellitus (T1DM) has been widely documented ^1–6^, but its pathophysiological origin remains poorly understood. For instance, between 1996 - 2015, T1DM incidence in Romanian children increased continuously at a rate that, if maintained, would double every 14 years ^7^. There have been numerous hypotheses presented in the literature to explain this rising incidence, many of which implicate aspects of the exposome. Notably, the ‘hygiene hypothesis’ for autoimmune and allergic diseases argues that factors affecting the skin or gut microbiome may be more important than infections ^8^ in the onset of T1DM in genetically predisposed individuals. On the other hand, Mistry et al. associated T1DM risk with sequences of infections ^9–13^ in genetically at-risk populations. Pathogenesis of T1DM has been also associated with the interplay between enterovirus and host microbiome ^14^. Moreover, there is a decreasing age at diagnosis of T1DM that could be accounted to the impact of environmental factors on children with lower-risk HLA class II genes, who previously would not have developed T1DM in childhood ^15^. Finally, there is no evidence of an association between routine vaccinations and childhood T1DM ^16^. These studies have been primarily limited to specific external exposures, and few studies examine aspects of the built environment in the rising incidence of T1DM.

After World War II, there was a great worldwide expansion of the plastics industry. According to author Susan Freinkel, “*In product after product, market after market, plastics challenged traditional materials and won, taking the place of steel in cars, paper and glass in packaging, and wood in furniture*” ^17^. In recent years, concerns about health effects of plastics, especially microplastics that have been found in multiple human tissues, have been reported ^18^ leading to the introduction of the concept of the human plastisphere ^19,20^.

Early literature, particularly in Scandinavia, shows that childhood T1DM was rare but well recognized before the introduction of insulin. Low incidence and prevalence rates continued over the period 1920 – 1950, with a carefully performed study showing no change in childhood incidence over the period 1925–1955 ^1^. As depicted in Figure 2 of Gale ^1^, the overall rate of T1DM diagnoses changed dramatically in the 1950s, shifting to a linear increase, with evidence of a plateau in some high-incidence populations and of a catch-up phenomenon in some low-incidence areas ^1^. To the authors knowledge, the wide presence of plastics in human samples ^19,20^ have not been associated with the increase of T1DM, but the obvious temporal correlation between the wide adoption of plastics and increase of T1DM in the 1950s is worthwhile considering.

Molecular mimicry as an autoimmune trigger ^21–25^ has long been recognized as a mechanism by which exogenous epitopes exhibiting similar sequence, structural, and/or binding properties to epitopes from disease antigens, can potentially trigger autoimmune diseases in genetically predisposed individuals. This has been recently studied in our group for T1DM ^26,27^ and for SARS-CoV-2 ^28^. Recent advances in protein structure prediction using artificial intelligence ^29–31^ allows us now to go beyond sequence homology and computationally explore binding of molecular fragments that do not have any homology with antigen epitopes. Using these emerging AI approaches, we can now attempt to answer the following: *Can short micro plastic fragments lead to pathogenesis of T1DM by molecular mimicry?* To explore this question, in this paper we report docking calculations of established T1DM antigen epitopes to respective HLA molecules and compare their docking with fragments of five commonly used plastics.

## Materials and Methods

Here we considered the HLA molecules corresponding to the most predisposing T1DM genotypes: *DRB1*\*03:01-*DQA1*\*05:01-*DQB1*\*02:01 and *DRB1*\*04:01/02/04/05/08-*DQA1*\*03:01-*DQB1*\*03:02/04 (or *DQB1*\*02) ^32^ and the HLA molecules identified in molecular mimicry by Mistry *et al*. ^33^ as potential binding substrates to induce molecular mimicry. The HLA molecules and respective T1DM antigen epitopes (between parenthesis) considered here are *HLA-*A*02:01 (RLLPLLALLAL), *HLA-*B*08:01 (TPKTRREAEDL); *HLA-DQA1*\*03:01 (FVNQHLCGSHLVEAL); *HLA-DQA1*\*05:01 (SHLVEALYLVCGERG), *HLA-DQB1*\*02:01 (SHLVEALYLVCGERG), *HLA-DQB1*\*03:02 (SHLVEALYLVCGERG), *HLA-DRB1*\*03:01 (LPKPPKPVSKMRMATPLLMQALPM), *HLA-DRB1*\*04:01 (NFFRMVISNPAAT), *HLA-* DRB3*02:02 (SPLGQSQPTVAGQPSARPAAEEYGYIVTDQKPLSLAAGVK), and *HLA-* DRB4*01:01 (KVNFFRMVISNPAATHQD) ^34^. The amino acid sequences of the HLA molecules were obtained from (https://www.ebi.ac.uk/ipd/imgt/hla/) and depicted in Table S1. The corresponding antigen epitopes were obtained from IEDB ^34^ (https://www.iedb.org/) and entered in Table S2.

Fragments of six common microplastics, high-density polyethylene (HDPE), low-density polyethylene (LDPE), polyethylene terephthalate (PET), polypropylene (PP), and polystyrene (PS) and polyvinyl chloride (PVC) were chosen as potential candidates for molecular mimicry. Their SMILES fragments ^35^ were retrieved from PubChem (https://pubchem.ncbi.nlm.nih.gov/) and were input into a python v.3.11.9 (with the RdKit library and AllChem module) script to generate covalently linked microplastic oligomers of desired length with corresponding structural data files (SDF). These SDF files were input into UCSF’s ChimeraX (v.1.10.1) ^36^ for structure analysis. The polymer length was measured using the ChimeraX ‘distance’ command. All six of the plastics fragments were selected to be of the approximate length of the binding region of the target Human Leukocyte Antigen (HLA) molecules. The SMILE notations for all the plastic fragments used here are entered into Figure S1.

Structural modeling of the HLA-T1DM epitope complexes and HLA-plastic fragments complexes were done using Boltz-2 (v.2.2.0), ^30^ an open-source AI affinity prediction model. Boltz 2 was executed on the Granite GPU cluster at the Utah Center for High Performance Computing (https://www.chpc.utah.edu/). Boltz-2 was run using standard parameters by enabling the MSA server, specifying 10 recycling steps, generating five diffusion samples, and performing 500 sampling steps. Each run generated 4 candidate models for each FASTA file (in CIF file format), but model 0 consistently exhibited the highest accuracy and highest confidence scores ^30^.

The output CIF files for each structural model were input into ChimeraX to visualize the calculated structure and to calculate structural similarity. All the Boltz-2 results are available at 10.5281/zenodo.19100817. The location of the T1DM epitopes and microplastic polymers were visually inspected to determine whether they resided in the binding region of HLA molecule and are depicted in Figure S2. The binding facts discussed under the results section of the paper have been extracted from the .cif files for all the models discussed here using the University of Utah instance of ChatGPT 5.2 on March 11, 2026 and the script used is given in the Supplementary Material. The ability of the plastic fragments to bind at the same groove as the T1DM epitope peptide was classified as Strong, Partial and No overlap using the following algorithm implemented in the ChatGPT 5.2. All results reported here were furthermore verified by one of the authors (JCF) and none of these results we directly imported from ChatGPT without human supervision, as in many cases the initial ChatGPT results were clearly erroneous and needed further prompting to clarify them. Clearly LLMs like ChatGPT can be used to analyze more efficiently .cif files, but currently the technology is not accurate enough to allow automatic analysis without expert supervision.

## Results

Among all the cases analyzed, model_0 from the Boltz-2 predictions consistently exhibited the highest confidence scores. For these models, the confidence scores show an average of 0.8090, with a standard deviation of 0.0480, a Max of 0.8979, and a Min of 0.6540. The scores for all models considered here are presented in the Supplementary Material Table S3. The great majority of these scores are considered very good ^30^, indicating that the results discussed here for all model structures are reliable computational predictions of the binding regions of antigen epitopes and plastic fragments to HLA molecules.

In the Supplementary Material Figures S2A-J we present a pictorial comparison of the predicted binding sites of the HLA molecule with corresponding T1DM antigen epitope and the six plastic fragments grouped by HLA haplotype. The corresponding .cif files can be found in the ZENODO repository: https://zenodo.org/records/19100817. For all the models, the structural features reported here have been extracted from the .cif files using the University of Utah instance of ChatGPT 5.2 and were furthermore verified by one of the authors (JCF). None of the results were directly imported from ChatGPT without human supervision.

We summarize the results in Table 1, where using the algorithm described in the Supplementary Material, we classified the plastic-HLA molecule interactions as showing strong, partial and no overlap in the binding grooves of the HLA-specific T1DM antigen epitopes. There are 19 systems showing strong overlaps, 17 showing partial overlaps, and 24 showing no overlaps. The plastic fragments are found in the following number models (Strong, Partial, No) as follows: HDPE (1,4,5), LDPE (3, 2, 5), PET (5,2, 3), PP (2, 5, 3), PS (4, 3, 3) and PVC (4, 1, 5) indicating that all of them are at least capable of inducing molecular mimicry with one of the haplotypes considered here. Moreover, non-HLA or plastic fragments show universal binding, arguing that the results presented here are not a consequence of wide binding promiscuity. Models showing Strong Overlap are displayed in Figure 1.

**Table 1:** Models considered in this study showing Strong, Partial and No Overlap with the Binding Location of the T1DM Antigen and the HLA molecule.

| **System** | **Strong Overlap** | **Partial Overlap** | **No Overlap** |
|---|---|---|---|
| HLA*A:02:01 | PS, PP, PET | PVC | HDPE, LDPE |
| HLA*B:08:01 | PS, PP, PET | — | PVC, HDPE, LDPE |
| HLA*DQA:03:01 | — | PET | HDPE, LDPE, PP, PS, PVC |
| HLA*DQA1:05:01 | PS | PET, HDPE | LDPE, PP, PVC |
| HLA*DQB1:02:01 | PET, LDPE, PVC | PP, PS, HDPE | — |
| HLA*DQB1:03:02 | PVC, PET, HDPE, LDPE | PP, PS | — |
| HLA*DRB1:03:01 | PET, LDPE | PP, PS | HDPE, PVC |
| HLA*DRB1:04:01 | PS, PVC | HDPE, PP, LDPE | PET |
| HLA*DRB3:02:02 | — | — | HDPE, LDPE, PET, PP, PS, PVC |
| HLA*DRB4:01:01 | PVC | HDPE, PP, LDPE | PET, PS |

**Figure 1.**
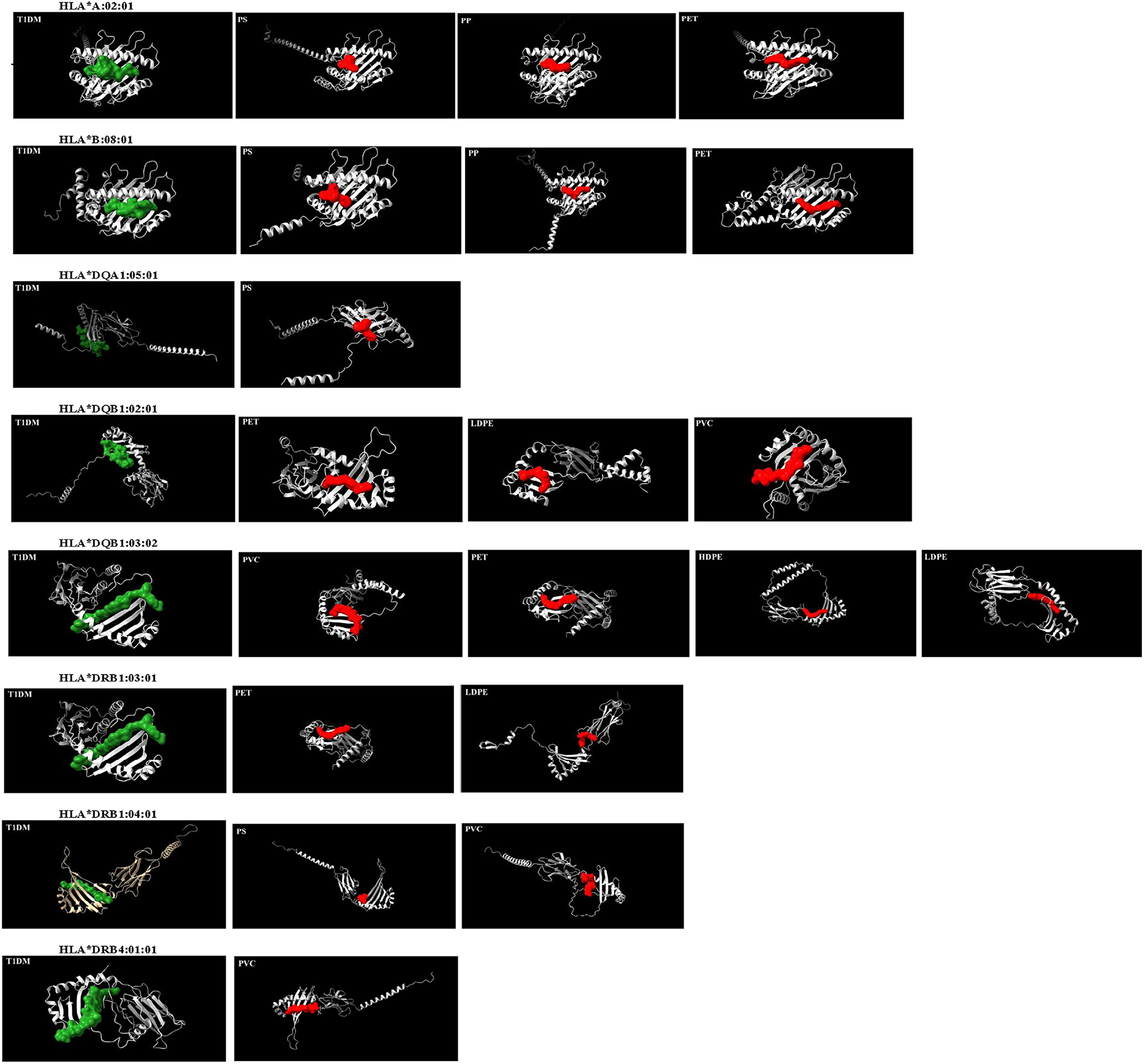
Models showing Strong Overlap in the binding grooves of HLA-specific T1DM antigen epitopes. The T1DM antigen epitopes are depicted in green, and the plastic fragments are depicted in red. Larger figures are available in the Supplementary Material and full .cif files can be found in the ZENODO repository https://zenodo.org/records/19100817.

In the models for HLA*A:02:01, Figure S2A, the T1DM antigen binds in an extended groove defined by the N-terminal region (residues ~31 to 105) and a C-terminal groove region (residues ~121 to 195). This is a classic MHC class II-like peptide binding cleft ^37^. The plastic fragments of PP, PS, and PET bind strongly in the same groove on the HLA molecule. The PVC fragment also partially binds in this region, while the HDPE and LDPE fragments, which are non-polar/hydrophobic, bind on the surface of the protein and are classified as having no overlap.

In the models for HLA*B:08:01, Figure S2B, the T1DM antigen binds in the same extended groove as in the HLA*A:02-01 models. Plastic fragments from PET, PP and PS bind in a similar region, showing significant overlap with the binding region of the T1DM antigen. The other fragments, HDPE, LDPE and PVC, bind in different regions.

In the models for HLA*DQA1:03:01, Figure S2C, the T1DM antigen binds in an extended surface groove roughly spanning residues ~16 to 102. In this case, none of the plastic fragments bind in a site that overlaps with the T1DM antigen binding site, but the PET fragment shows a modest overlap.

In the models for HLA*DQA1:05:01, Figure S2D, the T1DM antigen binds in a groove formed by residues roughly between ~30–90, with multiple clustered contact regions (30–37, 50–57, 69– 79, 83–87). This is a typical extended surface-binding mode in other DQA1/T1DM models ^38^.

The fragment of PS shows the strongest overlap with the T1DM peptide site (7 shared residues), suggesting it occupies the same groove and may compete with antigen peptide binding. PET and HDPE show partial overlaps (2 and 1 shared residues respectively) and they sit near or partially in the groove, while LDPE, PP, PVC bind distinct pockets with no meaningful overlap.

In the models for HLA*DQB1:02:01, Figure S2E, the T1DM antigen binds in a groove spanning residues ~41–117 with several contact clusters (around ~41–60 and ~89–117). This is consistent with an extended-peptide binding mode ^38^. Across these models, most plastic fragments overlap somehow with the peptide-binding site, particularly residues 41 and 43, which are common shared contacts across nearly all ligand models. PET, PVC and LDPE fragments show the strongest overlaps (4–5 shared residues), suggesting they occupy the peptide groove or a directly adjacent pocket, while PP, PS and HDPE fragments show only partial overlap.

In the models for HLA*DQB1:03:02, Figure S2F, the T1DM antigen binds in a groove covering residues roughly ~38–52 with additional contacts at 64, 115, and 123, mostly an N-terminal/central groove region. Many of these plastic fragments (especially PVC, PET, HDPE, LDPE, 4C) overlap substantially with the peptide-binding groove (notably residues around ~41– 50), suggesting they occupy or intrude into the peptide-binding site and could compete with peptide binding. The fragments of plastics PP and PS overlap less with the peptide site, indicating alternative pocket usage or partial peripheral interaction.

In the models for HLA*DRB1:03:01, Figure S2G, the T1DM antigen binds in an extended groove, and certain central residues (esp. positions ~10–11 and 18) make the most contact with the HLA molecule. Contact distribution (where several central residues with high contact counts flank residues with fewer contacts) is typical of peptide-binding grooves where a few peptide side chains sit in pockets and the backbone makes multiple interactions ^38^. Fragments of PET and LDPE show overlap with the peptide-binding groove (PET particularly overlaps residues ~42–45 which are central to the peptide interface). Also, fragments from PP and PS have some partial overlap, PP at the C-terminal peptide-contacting residues ~163/166 and PS touches residues ~115/182. Finally fragments from HDPE and PVC bind distinct pockets with no direct overlap to the peptide site.

In the models for HLA*DRB1:04:01, Figure S2H, the T1DM antigen binds to central residues (positions 2–6 and 4 especially), which act as primary anchors in this model. The contact pattern (clustered central anchors with multiple protein residues contacted) matches typical peptide-binding groove behavior ^38^. Several plastic fragments, notably PS and PVC and to a lesser extent the others, overlap with the peptide-binding groove (shared residues include 46, 111, and 119 frequently). The PET fragment does not show any overlapping.

In the models for HLA*DRB3:02:02, Figure S2I, the T1DM antigen binds with multiple surface clusters on chain B (examples include residues around ~120–130, ~170–185, and several in the ~20–50 region). Positions ~25 and ~30 appear to be the strongest local anchors in this model. None of the six plastic fragments show significant overlap with the peptide-contact residues. In other words, the modeled ligands occupy pockets/surface patches distinct from the peptide interface in this HLA*DRB3:02:02 model set.

In the models for HLA*DRB4:01:01, Figure S2J, the T1DM antigen binds in an extended groove on chain B with a dense cluster of contacts around residues ~35–46 and another region at ~262– 266, indicating central anchor positions in the 4–7 range and additional stabilization at the C-terminal end. The plastic fragment of PVC shows the largest intrusion into the peptide groove (4 shared residues), followed by HDPE, LDPE, and PP with small overlaps (1–2 residues), indicating that they could compete with peptide binding. Fragments of PET and PS bind distinct pockets with no overlap to the peptide groove.

Even when considering the worst-case scenario, in which only the plastic fragments that show strong overlap (19 cases found here) with the binding location of the T1DM antigen and the HLA molecule, there are significant opportunities for multiple plastic fragments to potentially induce molecular mimicry establishing plausible pathophysiological mechanisms for which plastics could trigger T1DM in individuals with certain HLA haplotypes.

## Conclusions

Recent advances in artificial intelligence for protein structure and binding modeling, like Boltz-2 ^30,31^, allows researchers to computationally explore hypotheses that may not be amenable to experimental or epidemiological studies. Here we show that Boltz-2 can predict T1DM antigen binding consistent with previous experimental work ^37,38^. The models showing binding of plastic polymers, for which experimental studies are not available, show similar levels of confidence to those from the T1DM antigens. The results here show that plastic fragments that can be found in many human organs ^18,20^, can bind to HLA molecules at the same sites that well known epitopes from T1DM antigens bind, leading to potential molecular mimicry, similar to how infectious epitopes can trigger T1DM and other autoimmune diseases ^9,28,33^. While the results presented here do not prove causality between potential molecular mimicry by plastic fragments, the results provide the first pathophysiological hypothesis of a plausible molecular mechanism that explains the temporal correlation on the significant increase of the onset of T1DM and the expanded use of plastics in the 1950s. While the results suggest that most common plastics can bind to at least one of the HLA sites recognized by T1DM epitopes across the six HLA molecules analyzed, the limited scope of HLA alleles examined prevents any strong conclusions about which specific haplotypes may be more prone to binding plastic fragments.

## Supporting information

All suplementary material

## Acknowledgments

Computational resources were provided by the Utah Center for High Performance Computing. Molecular graphics and analyses were performed with UCSF ChimeraX, developed by the Resource for Biocomputing, Visualization, and Informatics at the University of California, San Francisco, with support from National Institutes of Health R01-GM129325 and the Office of Cyber Infrastructure and Computational Biology, National Institute of Allergy and Infectious Diseases.

## Author contributions

All authors contributed to the formulation of the problem. MJ and JCF performed the Boltz-2 calculations. JCF wrote the first draft of the paper and all authors revised and approved the final version of the paper

## Data, code, and materials availability

Boltz-2 is an open-source software available at: https://github.com/jwohlwend/boltz

All the FASTA files and Boltz-2 results are available at ZENDO 10.5281/zenodo.19100817

## Statement on use of AI

The binding facts discussed under the results section of the paper have been extracted from the .cif files for all the models discussed here using the University of Utah instance of ChatGPT 5.2 on March 11, 2026. The script used is given in the Supplementary Material. All results reported here were furthermore verified by one of the authors (JCF) and none of these results we directly imported from ChatGPT without human supervision, as in many cases the initial ChatGPT results were clearly erroneous and needed further prompting to clarify them.

## References and Notes

1. Gale EAM. The Rise of Childhood Type 1 Diabetes in the 20th Century. Diabetes. 2002;51(12):3353–3361. doi:10.2337/diabetes.51.12.3353

2. Bandarian F, Sharifnejad Tehrani Y, Peimani M, et al. National and sub-national burden and trend of type 1 diabetes in 31 provinces of Iran, 1990-2019. Sci Rep. Mar 14 2023;13(1):4210. doi:10.1038/s41598-023-31096-8

3. Dabelea D, Sauder KA, Jensen ET, et al. Twenty years of pediatric diabetes surveillance: what do we know and why it matters. Ann N Y Acad Sci. Jul 2021;1495(1):99–120. doi:10.1111/nyas.14573

4. Evertsen J, Alemzadeh R, Wang X. Increasing incidence of pediatric type 1 diabetes mellitus in Southeastern Wisconsin: relationship with body weight at diagnosis. PLoS One. Sep 3 2009;4(9):e6873. doi:10.1371/journal.pone.0006873

5. Niechciał E, Michalak M, Skowrońska B, Fichna P. Increasing trend of childhood type 1 diabetes incidence: 20-year observation from Greater Poland Province, Poland. Acta Diabetol. Dec 2024;61(12):1609–1617. doi:10.1007/s00592-024-02339-5

6. Norris JM, Johnson RK, Stene LC. Type 1 diabetes—early life origins and changing epidemiology. The Lancet Diabetes & Endocrinology. 2020;8(3):226–238. doi:10.1016/S2213-8587(19)30412-7

7. Vlad A, Serban V, Green A, et al. Time Trends, Regional Variability and Seasonality Regarding the Incidence of Type 1 Diabetes Mellitus in Romanian Children Aged 0-14 Years, Between 1996 and 2015. J Clin Res Pediatr Endocrinol. Jun 1 2018;10(2):92–99. doi:10.4274/jcrpe.5456

8. Abela AG, Fava S. Prenatal and early life factors and type 1 diabetes. Endocrine. Jun 2022;77(1):48–56. doi:10.1007/s12020-022-03057-0

9. Mistry S, Gouripeddi R, Raman V, Facelli JC. Sequential data mining of infection patterns as predictors for onset of type 1 diabetes in genetically at-risk individuals. Journal of Biomedical Informatics. 2023:104385.

10. Bougnères P, Le Fur S, Valtat S, Kamatani Y, Lathrop M, Valleron AJ. Using spatiotemporal surveillance data to test the infectious environment of children before type 1 diabetes diagnosis. PLoS One. 2017;12(2):e0170658. doi:10.1371/journal.pone.0170658

11. Dedrick S, Sundaresh B, Huang Q, et al. The Role of Gut Microbiota and Environmental Factors in Type 1 Diabetes Pathogenesis. Front Endocrinol (Lausanne). 2020;11:78. doi:10.3389/fendo.2020.00078

12. Fenneman AC, Weidner M, Chen LA, Nieuwdorp M, Blaser MJ. Antibiotics in the pathogenesis of diabetes and inflammatory diseases of the gastrointestinal tract. Nat Rev Gastroenterol Hepatol. Feb 2023;20(2):81–100. doi:10.1038/s41575-022-00685-9

13. Nishioka Y, Noda T, Okada S, et al. Association between influenza and the incidence rate of new-onset type 1 diabetes in Japan. J Diabetes Investig. Oct 2021;12(10):1797–1804. doi:10.1111/jdi.13540

14. Hober D, Sauter P. Pathogenesis of type 1 diabetes mellitus: interplay between enterovirus and host. Nat Rev Endocrinol. May 2010;6(5):279–89. doi:10.1038/nrendo.2010.27

15. Fourlanos S, Varney MD, Tait BD, et al. The rising incidence of type 1 diabetes is accounted for by cases with lower-risk human leukocyte antigen genotypes. Diabetes Care. Aug 2008;31(8):1546–9. doi:10.2337/dc08-0239

16. Morgan E, Halliday SR, Campbell GR, Cardwell CR, Patterson CC. Vaccinations and childhood type 1 diabetes mellitus: a meta-analysis of observational studies. Diabetologia. Feb 2016;59(2):237–43. doi:10.1007/s00125-015-3800-8

17. Freinkel S. Plastics: A Toxic Love Story (Henry Holt; 2011.

18. Evode N, Qamar SA, Bilal M, Barceló D, Iqbal HMN. Plastic waste and its management strategies for environmental sustainability. Case Studies in Chemical and Environmental Engineering. 2021/12/01/2021;4:100142. doi:10.1016/j.cscee.2021.100142

19. Hegde S, Min KT, Moore J, et al. Indoor Household Particulate Matter Measurements Using a Network of Low-cost Sensors. Aerosol and Air Quality Research. 2020;20(2):381–394. doi:10.4209/aaqr.2019.01.0046

20. Ziani K, Ioniţă-Mîndrican CB, Mititelu M, et al. Microplastics: A Real Global Threat for Environment and Food Safety: A State of the Art Review. Nutrients. Jan 25 2023;15(3) doi:10.3390/nu15030617

21. Rojas M, Herrán M, Ramírez-Santana C, et al. Molecular mimicry and autoimmunity in the time of COVID-19. Journal of Autoimmunity. 2023/09/01/2023;139:103070. doi:10.1016/j.jaut.2023.103070

22. Rojas M, Restrepo-Jiménez P, Monsalve DM, et al. Molecular mimicry and autoimmunity. Journal of Autoimmunity. 2018/12/01/2018;95:100–123. doi:10.1016/j.jaut.2018.10.012

23. Coppieters KT, Wiberg A, von Herrath MG. Viral infections and molecular mimicry in type 1 diabetes. Apmis. Dec 2012;120(12):941–9. doi:10.1111/apm.12011

24. Fujinami RS, von Herrath MG, Christen U, Whitton JL. Molecular mimicry, bystander activation, or viral persistence: infections and autoimmune disease. Clin Microbiol Rev. Jan 2006;19(1):80–94. doi:10.1128/cmr.19.1.80-94.2006

25. Peterson LK, Fujinami RS. 3 - MOLECULAR MIMICRY. In: Shoenfeld Y, Gershwin ME, Meroni PL, eds. Autoantibodies (Second Edition). Elsevier; 2007:13–19.

26. Mistry S, Gouripeddi R, Facelli JC. Prioritization of infectious epitopes for translational investigation in type 1 diabetes etiology. Article. Journal of Autoimmunity. 2023;140103115. doi:10.1016/j.jaut.2023.103115

27. Gardner R, Wilkins J, Mistry S, Gouripeddi R, Facelli JC. Structural Homology of Epitope Binding Mimicry in the Onset of Type 1 Diabetes Mellitus. IEEE; 2023:3708–3713.

28. Maldonado-Catala P, Gouripeddi R, Schlesinger N, Facelli JC. Molecular mimicry impact of the COVID-19 pandemic: Sequence homology between SARS-CoV-2 and autoimmune diseases epitopes. ImmunoInformatics. 2025;18:100050.

29. Varadi M, Anyango S, Deshpande M, et al. AlphaFold Protein Structure Database: massively expanding the structural coverage of protein-sequence space with high-accuracy models. Nucleic Acids Research. 2022;50(D1):D439–D444. doi:10.1093/nar/gkab1061

30. Passaro S, Corso G, Wohlwend J, et al. Boltz-2: Towards Accurate and Efficient Binding Affinity Prediction. bioRxiv. 2025:2025.06.14.659707. doi:10.1101/2025.06.14.659707

31. Wohlwend J, Corso G, Passaro S, et al. Boltz-1 Democratizing Biomolecular Interaction Modeling. bioRxiv. May 6 2025; doi:10.1101/2024.11.19.624167

32. Noble JA, Valdes AM. Genetics of the HLA region in the prediction of type 1 diabetes. Curr Diab Rep. Dec 2011;11(6):533–42. doi:10.1007/s11892-011-0223-x

33. Mistry S, Gouripeddi R, Facelli JC. Prioritization of infectious epitopes for translational investigation in type 1 diabetes etiology. Journal of Autoimmunity. 2023;140:103115.

34. Vita R, Mahajan S, Overton JA, et al. The Immune Epitope Database (IEDB): 2018 update. Nucleic Acids Res. Jan 8 2019;47(D1):D339–d343. doi:10.1093/nar/gky1006

35. Weininger D, Weininger A, Weininger JL. SMILES. 2. Algorithm for generation of unique SMILES notation. Journal of Chemical Information and Computer Sciences. 1989/05/01 1989;29(2):97–101. doi:10.1021/ci00062a008

36. Pettersen EF, Goddard TD, Huang CC, et al. UCSF ChimeraX: Structure visualization for researchers, educators, and developers. Protein Sci. Jan 2021;30(1):70–82. doi:10.1002/pro.3943

37. Bjorkman PJ, Saper MA, Samraoui B, Bennett WS, Strominger JL, Wiley DC. Structure of the human class I histocompatibility antigen, HLA-A2. Nature. 1987/10/01 1987;329(6139):506–512. doi:10.1038/329506a0

38. Lee KH, Wucherpfennig KW, Wiley DC. Structure of a human insulin peptide-HLA-DQ8 complex and susceptibility to type 1 diabetes. Nat Immunol. Jun 2001;2(6):501–7. doi:10.1038/88694

