## Supplementary material for "Can the Human Plastisphere Account for the Rise of Type 1 Diabetes: A Computational Molecular Mimicry Study": All suplementary material

**ChatGPT script**

Define the peptide-binding site

From each T1DM model:

Identify all protein residues (chain B) within ≤ 4.0 Å of the peptide (chain

👉 This gives the:

Peptide-binding footprint (groove residues)

________________________________________

2️⃣ Define ligand binding site

From each ligand model:

Identify all protein residues within ≤ 4.0 Å of the ligand

👉 This gives the:

Ligand-contact footprint

________________________________________

3️⃣ Compute overlap

Compare the two sets:

"Overlap"="Peptide residues"∩"Ligand residues"

________________________________________

📊 Classification thresholds (what defines “strong”, “partial”, “none”)

This is the key part 👇

🟢 Strong overlap (same groove)

≥ ~40–50% of ligand-contact residues overlap

OR

Ligand contacts core groove residues (central cluster)

👉 Interpretation:

Ligand is sitting in the peptide-binding cleft

Likely direct competition

________________________________________

🟡 Partial overlap

~10–40% overlap

Often only edge residues or one cluster overlaps

👉 Interpretation:

Ligand is near or partially intruding into groove

Possible weak or indirect competition

________________________________________

🔵 No overlap

0% overlap

No shared residues

👉 Interpretation:

Ligand binds a completely different pocket

No direct competition

________________________________________

**Table S1.** FASTA files for HLA molecules used as input into Boltz-2 for biomolecular modeling (https://www.ebi.ac.uk/ipd/imgt/hla/).

**HLA-A*02:01**

MAVMAPRTLVLLLSGALALTQTWAGSHSMRYFFTSVSRPGRGEPRFIAVGYVDDTQFVRFDSDAASQRMEPRAPWIEQEGPEYWDGETRKVKAHSQTHRVDLGTLRGYYNQSEAGSHTVQRMYGCDVGSDWRFLRGYHQYAYDGKDYIALKEDLRSWTAADMAAQTTKHKWEAAHVAEQLRAYLEGTCVEWLRRYLENGKETLQRTDAPKTHMTHHAVSDHEATLRCWALSFYPAEITLTWQRDGEDQTQDTELVETRPAGDGTFQKWAAVVVPSGQEQRYTCHVQHEGLPKPLTLRWEPSSQPTIPIVGIIAGLVLFGAVITGAVVAAVMWRRKSSDRKGGSYSQAASSDSAQGSDVSLTACKV

**HLA-B*08:01**

MLVMAPRTVLLLLSAALALTETWAGSHSMRYFDTAMSRPGRGEPRFISVGYVDDTQFVRFDSDAASPREEPRAPWIEQEGPEYWDRNTQIFKTNTQTDRESLRNLRGYYNQSEAGSHTLQSMYGCDVGPDGRLLRGHNQYAYDGKDYIALNEDLRSWTAADTAAQITQRKWEAARVAEQDRAYLEGTCVEWLRRYLENGKDTLERADPPKTHVTHHPISDHEATLRCWALGFYPAEITLTWQRDGEDQTQDTELVETRPAGDRTFQKWAAVVVPSGEEQRYTCHVQHEGLPKPLTLRWEPSSQSTVPIVGIVAGLAVLAVVVIGAVVAAVMCRRKSSGGKGGSYSQAACSDSAQGSDVSLTA

***HLA-DQA1**03:01**

MILNKALMLGALALTTVMSPCGGEDIVADHVASYGVNLYQSYGPSGQYSHEFDGDEEFYVDLERKETVWQLPLFRRFRRFDPQFALTNIAVLKHNLNIVIKRSNSTAATNEVPEVTVFSKSPVTLGQPNTLICLVDNIFPPVVNITWLSNGHSVTEGVSETSFLSKSDHSFFKISYLTFLPSADEIYDCKVEHWGLDEPLLKHWEPEIPTPMSELTETVVCALGLSVGLVGIVVGTVLIIRGLRSVGASRHQGPL

***HLA-DQA1**05:01**

MILNKALMLGALALTTVMSPCGGEDIVADHVASYGVNLYQSYGPSGQYTHEFDGDEQFYVDLGRKETVWCLPVLRQFRFDPQFALTNIAVLKHNLNSLIKRSNSTAATNEVPEVTVFSKSPVTLGQPNILICLVDNIFPPVVNITWLSNGHSVTEGVSETSFLSKSDHSFFKISYLTLLPSAEESYDCKVEHWGLDKPLLKHWEPEIPAPMSELTETVVCALGLSVGLVGIVVGTVFIIRGLRSVGASRHQGPL

***HLA-DQB1**02:01**

MSWKKALRIPGGLRAATVTLMLSMLSTPVAEGRDSPEDFVYQFKGMCYFTNGTERVRLVSRSIYNREEIVRFDSDVGEFRAVTLLGLPAAEYWNSQKDILERKRAAVDRVCRHNYQLELRTTLQRRVEPTVTISPSRTEALNHHNLLVCSVTDFYPAQIKVRWFRNDQEETAGVVSTPLIRNGDWTFQILVMLEMTPQRGDVYTCHVEHPSLQSPITVEWRAQSESAQSKMLSGIGGFVLGLIFLGLGLIIHHRSQKGLLH

***HLA-DQB1**03:02**

MSWKKALRIPGGLRVATVTLMLAMLSTPVAEGRDSPEDFVYQFKGMCYFTNGTERVRLVTRYIYNREEYARFDSDVGVYRAVTPLGPPAAEYWNSQKEVLERTRAELDTVCRHNYQLELRTTLQRRVEPTVTISPSRTEALNHHNLLVCSVTDFYPAQIKVRWFRNDQEETTGVVSTPLIRNGDWTFQILVMLEMTPQRGDVYTCHVEHPSLQNPIIVEWRAQSESAQSKMLSGIGGFVLGLIFLGLGLIIHHRSQKGLLH

***HLA-DRB1**03:01**

MVCLRLPGGSCMAVLTVTLMVLSSPLALAGDTRPRFLEYSTSECHFFNGTERVRYLDRYFHNQEENVRFDSDVGEFRAVTELGRPDAEYWNSQKDLLEQKRGRVDNYCRHNYGVVESFTVQRRVHPKVTVYPSKTQPLQHHNLLVCSVSGFYPGSIEVRWFRNGQEEKTGVVSTGLIHNGDWTFQTLVMLETVPRSGEVYTCQVEHPSVTSPLTVEWRARSESAQSKMLSGVGGFVLGLLFLGAGLFIYFRNQKGHSGLQPRGFLS

***HLA-DRB1**04:01**

MVCLKFPGGSCMAALTVTLMVLSSPLALAGDTRPRFLEQVKHECHFFNGTERVRFLDRYFYHQEEYVRFDSDVGEYRAVTELGRPDAEYWNSQKDLLEQKRAAVDTYCRHNYGVGESFTVQRRVYPEVTVYPAKTQPLQHHNLLVCSVNGFYPGSIEVRWFRNGQEEKTGVVSTGLIQNGDWTFQTLVMLETVPRSGEVYTCQVEHPSLTSPLTVEWRARSESAQSKMLSGVGGFVLGLLFLGAGLFIYFRNQKGHSGLQPTGFLS

***HLA-*DRB3*02:02**

MVCLKLPGGSSLAALTVTLMVLSSRLAFAGDTRPRFLELLKSECHFFNGTERVRFLERHFHNQEEYARFDSDVGEYRAVRELGRPDAEYWNSQKDLLEQKRGQVDNYCRHNYGVGESFTVQRRVHPQVTVYPAKTQPLQHHNLLVCSVSGFYPGSIEVRWFRNGQEEKAGVVSTGLIQNGDWTFQTLVMLETVPRSGEVYTCQVEHPSVTSPLTVEWSARSESAQSKMLSGVGGFVLGLLFLGAGLFIYFRNQKGHSGLQPTGFLS

***HLA-*DRB4*01:01**

MVCLKLPGGSCMAALTVTLTVLSSPLALAGDTQPRFLEQAKCECHFLNGTERVWNLIRYIYNQEEYARYNSDLGEYQAVTELGRPDAEYWNSQKDLLERRRAEVDTYCRYNYGVVESFTVQRRVQPKVTVYPSKTQPLQHHNLLVCSVNGFYPGSIEVRWFRNSQEEKAGVVSTGLIQNGDWTFQTLVMLETVPRSGEVYTCQVEHPSMMSPLTVQWSARSESAQSKMLSGVGGFVLGLLFLGTGLFIYFRNQKGHSGLQPTGLLS

**Table S2.** Epitopes associated with the HLA Molecules are considered here.

***HLA-*A*02:01**

RLLPLLALLAL: A linear peptidic epitope (epitope ID 115471) studied as part of Insulin (UniProt:P01308) from Homo sapiens (human). This epitope has been studied for immune reactivity in two publication(s), tested in four T cell assays and two MHC ligand assays.

***HLA-*B*08:01**

TPKTRREAEDL**:** A linear peptidic epitope (epitope ID 106918) studied as part of Insulin (UniProt:P01308) from Homo sapiens (human). This epitope has been studied for immune reactivity in one publication(s), tested in three T cell assays and one MHC ligand assay.

***HLA-DQA1**03:01**

FVNQHLCGSHLVEAL: A linear peptidic epitope (epitope ID 18247) studied as part of Insulin, isoform 2 (UniProt:F8WCM5) from Homo sapiens (human) and Insulin (UniProt:P01308) from Homo sapiens (human). This epitope has been studied for immune reactivity in seven publication(s), tested in seven T cell assays and 36 MHC ligand assays*.*

***HLA-DQA1**05:01**

SHLVEALYLVCGERG: A linear peptidic epitope (epitope ID 58388) studied as part of Insulin-2 (UniProt:P01326) from Mus musculus (mouse), Insulin (UniProt:P01308) from Homo sapiens (human) and Insulin, isoform 2 (UniProt:F8WCM5) from Homo sapiens (human). This epitope has been studied for immune reactivity in 79 publication(s), tested in 238 T cell assays, 31 B cell assays, 26 MHC ligand assays and has 3D structure(s) 1JK8

***HLA-DQB1**02:01**

SHLVEALYLVCGERG: A linear peptidic epitope (epitope ID 58388) studied as part of Insulin-2 (UniProt:P01326) from Mus musculus (mouse), Insulin (UniProt:P01308) from Homo sapiens (human) and Insulin, isoform 2 (UniProt:F8WCM5) from Homo sapiens (human). This epitope has been studied for immune reactivity in 79 publication(s), tested in 238 T cell assays, 31 B cell assays, 26 MHC ligand assays and has 3D structure(s) 1JK8.

***HLA-DQB1**03:02**

SHLVEALYLVCGERG; A linear peptidic epitope (epitope ID 58388) studied as part of Insulin-2 (UniProt:P01326) from Mus musculus (mouse), Insulin (UniProt:P01308) from Homo sapiens (human) and Insulin, isoform 2 (UniProt:F8WCM5) from Homo sapiens (human). This epitope has been studied for immune reactivity in 79 publication(s), tested in 238 T cell assays, 31 B cell assays, 26 MHC ligand assays and has 3D structure(s) 1JK8.

***HLA-DRB1**03:01**

LPKPPKPVSKMRMATPLLMQALPM: A linear peptidic epitope (epitope ID 38564) studied as part of HLA class II histocompatibility antigen gamma chain (UniProt:P04233) from Homo sapiens (human). This epitope has been studied for immune reactivity in 36 publication(s), tested in two T cell assays, 122 MHC ligand assays and has 3D structure(s) 1MUJ and 1A6A*.*

***HLA-DRB1**04:01**

NFFRMVISNPAAT: A linear peptidic epitope (epitope ID 101067) studied as part of Glutamate decarboxylase 2 (UniProt:Q05329) from Homo sapiens (human), Glutamate decarboxylase 2 (UniProt:P48320) from Mus musculus (mouse) and Glutamate decarboxylase 1 (UniProt:Q99259) from Homo sapiens (human). This epitope has been studied for immune reactivity in 23 publication(s), tested in 85 T cell assays, six B cell assays and five MHC ligand assays.

***HLA-*DRB3*02:02**

**S**PLGQSQPTVAGQPSARPAAEEYGYIVTDQKPLSLAAGVK**:** A linear peptidic epitope (epitope ID 534789) studied as part of Receptor-type tyrosine-protein phosphatase-like N (UniProt:Q16849) from Homo sapiens (human). This epitope has been studied for immune reactivity in one publication(s), tested in four T cell assays and one MHC ligand assay.

***HLA-*DRB4*01:01**

KVNFFRMVISNPAATHQD**:** A linear peptidic epitope (epitope ID 102589) studied as part of Glutamate decarboxylase 2 (UniProt:Q05329) from Homo sapiens (human). This epitope has been studied for immune reactivity in one publication(s), tested in one T cell assay and 2 MHC ligand assays.

**Table S3.** Boltz-2 confidence scores for all the models used in this study.

| **HLA Molecule** | **Confidence Score** |
| --- | --- |
| HLA-A-02-01_HDPE | 0.897938371 |
| HLA-A-02-01_LDPE | 0.867696941 |
| HLA-A-02-01_PET | 0.858108997 |
| HLA-A-02-01_PP | 0.871602237 |
| HLA-A-02-01_PS | 0.860020757 |
| HLA-A-02-01_PVC | 0.878630936 |
| HLA-A-02-01_T1D | 0.869014919 |
| HLA-B-08-01_HDPE | 0.849690557 |
| HLA-B-08-01_LDPE | 0.850920677 |
| HLA-B-08-01_PET | 0.816228032 |
| HLA-B-08-01_PP | 0.866624177 |
| HLA-B-08-01_PS | 0.865987778 |
| HLA-B-08-01_PVC | 0.774102092 |
| HLA-B-08-01_T1D | 0.856066525 |
| HLA-DQA1-03-01_HDPE | 0.849064946 |
| HLA-DQA1-03-01_LDPE | 0.830928922 |
| HLA-DQA1-03-01_PET | 0.791678488 |
| HLA-DQA1-03-01_PP | 0.852097034 |
| HLA-DQA1-03-01_PS | 0.791954815 |
| HLA-DQA1-03-01_PVC | 0.819305897 |
| HLA-DQA1-03-01_T1D | 0.816486657 |
| HLA-DQA1-05-01_HDPE | 0.834633648 |
| HLA-DQA1-05-01_LDPE | 0.840355992 |
| HLA-DQA1-05-01_PET | 0.797511876 |
| HLA-DQA1-05-01_PP | 0.829351068 |
| HLA-DQA1-05-01_PS | 0.830265224 |
| HLA-DQA1-05-01_PVC | 0.782985926 |
| HLA-DQA1-05-01_T1D | 0.855341852 |
| HLA-DQB1-02-01_HDPE | 0.848351657 |
| HLA-DQB1-02-01_LDPE | 0.8387025 |
| HLA-DQB1-02-01_PET | 0.766355932 |
| HLA-DQB1-02-01_PP | 0.869483113 |
| HLA-DQB1-02-01_PS | 0.77932477 |
| HLA-DQB1-02-01_PVC | 0.791695118 |
| HLA-DQB1-02-01_T1D | 0.870742917 |
| HLA-DQB1-03-02_HDPE | 0.806219041 |
| HLA-DQB1-03-02_LDPE | 0.809415638 |
| HLA-DQB1-03-02_PET | 0.817177236 |
| HLA-DQB1-03-02_PP | 0.80591327 |
| HLA-DQB1-03-02_PS | 0.827299297 |
| HLA-DQB1-03-02_PVC | 0.784492075 |
| HLA-DQB1-03-02_T1D | 0.817025006 |
| HLA-DRB1-03-01_HDPE | 0.749644578 |
| HLA-DRB1-03-01_LDPE | 0.767179787 |
| HLA-DRB1-03-01_PET | 0.791909516 |
| HLA-DRB1-03-01_PP | 0.741444767 |
| HLA-DRB1-03-01_PS | 0.77523309 |
| HLA-DRB1-03-01_PVC | 0.718434513 |
| HLA-DRB1-03-01_T1D | 0.727432072 |
| HLA-DRB1-04-01_HDPE | 0.804182827 |
| HLA-DRB1-04-01_LDPE | 0.834283054 |
| HLA-DRB1-04-01_PET | 0.786955953 |
| HLA-DRB1-04-01_PP | 0.826202512 |
| HLA-DRB1-04-01_PS | 0.774005115 |
| HLA-DRB1-04-01_PVC | 0.761941612 |
| HLA-DRB1-04-01_T1D | 0.781782269 |
| HLA-DRB3-02-02_HDPE | 0.822908342 |
| HLA-DRB3-02-02_LDPE | 0.849228084 |
| HLA-DRB3-02-02_PET | 0.775658071 |
| HLA-DRB3-02-02_PP | 0.870815694 |
| HLA-DRB3-02-02_PS | 0.84092325 |
| HLA-DRB3-02-02_PVC | 0.813858032 |
| HLA-DRB3-02-02_T1D | 0.654037893 |
| HLA-DRB4-01-01_HDPE | 0.762024403 |
| HLA-DRB4-01-01_LDPE | 0.770341218 |
| HLA-DRB4-01-01_PET | 0.739925086 |
| HLA-DRB4-01-01_PP | 0.751816511 |
| HLA-DRB4-01-01_PS | 0.719406247 |
| HLA-DRB4-01-01_PVC | 0.731726825 |
| HLA-DRB4-01-01_T1D | 0.751129746 |

**Figure S1.** SMILE notation and structures of the plastic fragments used in this study (https://www.leskoff.com/s01810-0).

**High-Density Polyethylene (HDPE)**

CCCCCCCCCCCCCCCCCCCC

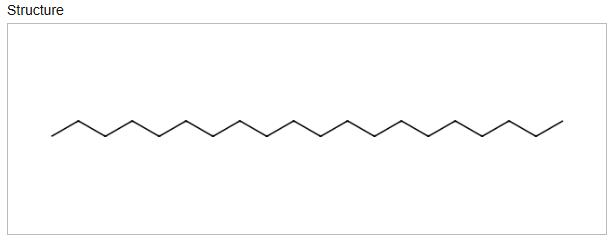

**Low-Density Polyethylene	(LDPE)**

CCCCCCC(C)CCCC(C)CCCCCC(CC)CCCC

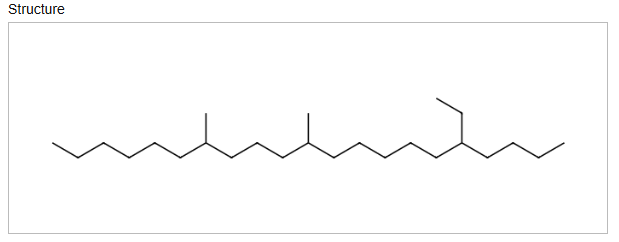

**Polyethylene Terephthalate (PET)**

O=C(c1ccc(C(=O)OCCO)cc1)OCCOOC(c1ccc(C(=O)OCCO)cc1)O

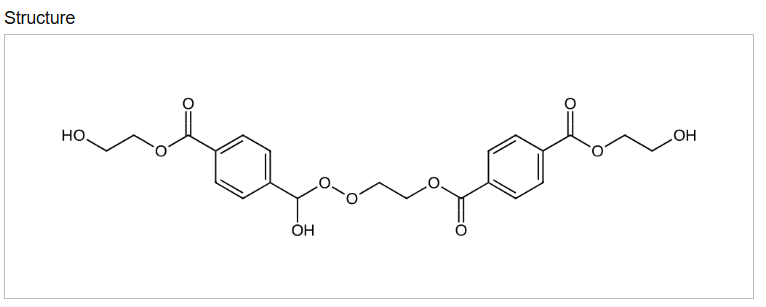

**Polypropylene (PP)**

CC(C)CC(C)CC(C)CC(C)CC(C)CC(C)CC(C)CC(C)CC

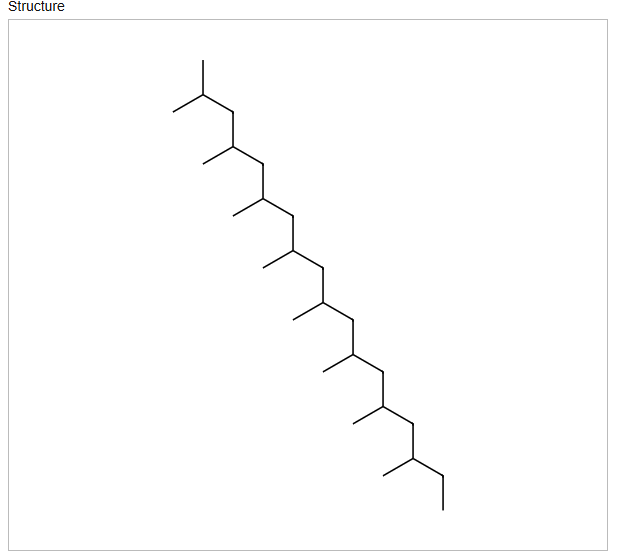

**Polystyrene (PS)**

CC(C1=CC=CC=C1)CC(C1=CC=CC=C1)CC(C1=CC=CC=C1)CC(C1=CC=CC=C1)

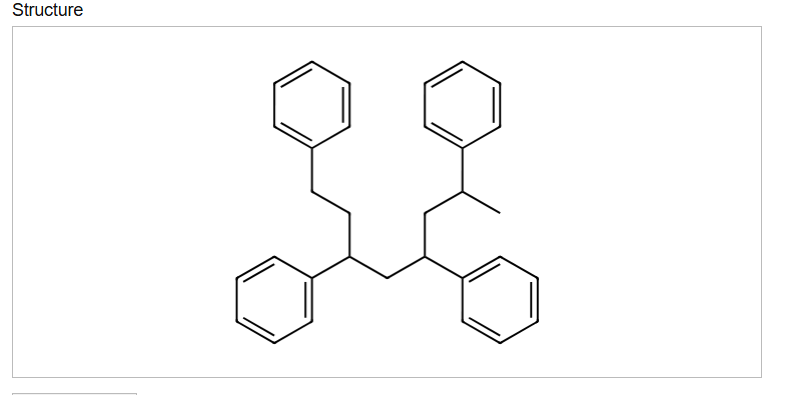

**Polyvinyl Chloride (PVC)**

CC(Cl)CC(Cl)CC(Cl)CC(Cl)CC(Cl)CC(Cl)CC(Cl)CC(Cl)CC(Cl)CC(Cl)CC(Cl)CC(Cl)CC(Cl)

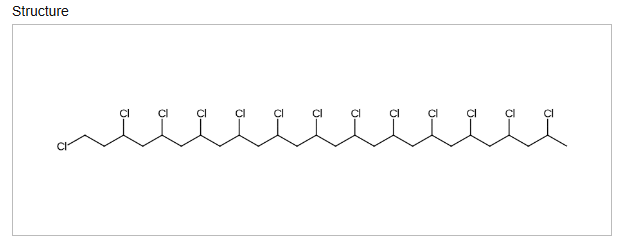

**Figure S2A**: Model structures of all HLA-A*02:01 systems studied here. The T1DM antigen epitopes are depicted in green, and the plastic fragments are depicted in red.

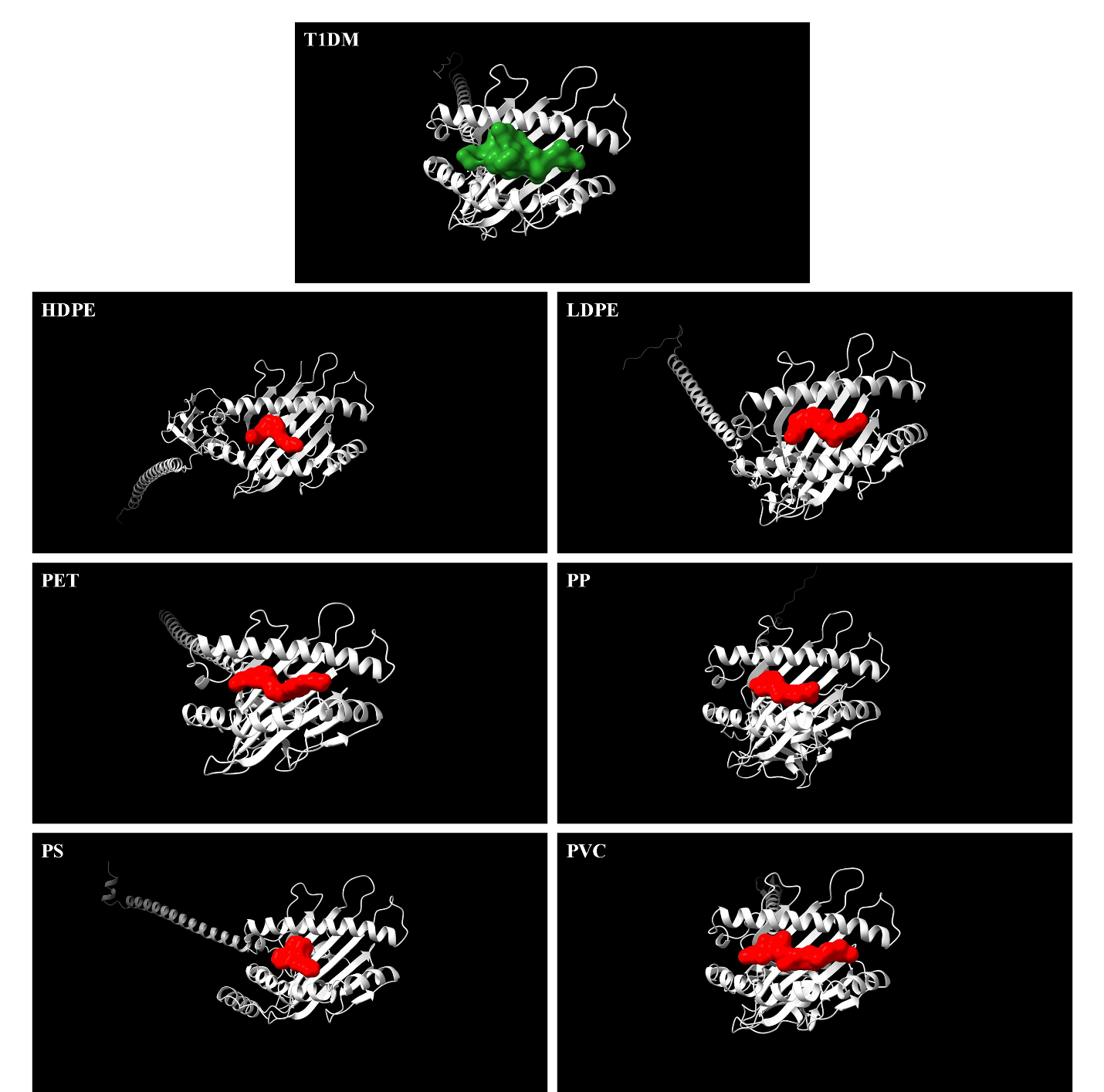

**Figure S2B**: Model structures of all HLA-B*08:01 systems studied here. The T1DM antigen epitopes are depicted in green, and the plastic fragments are depicted in red.

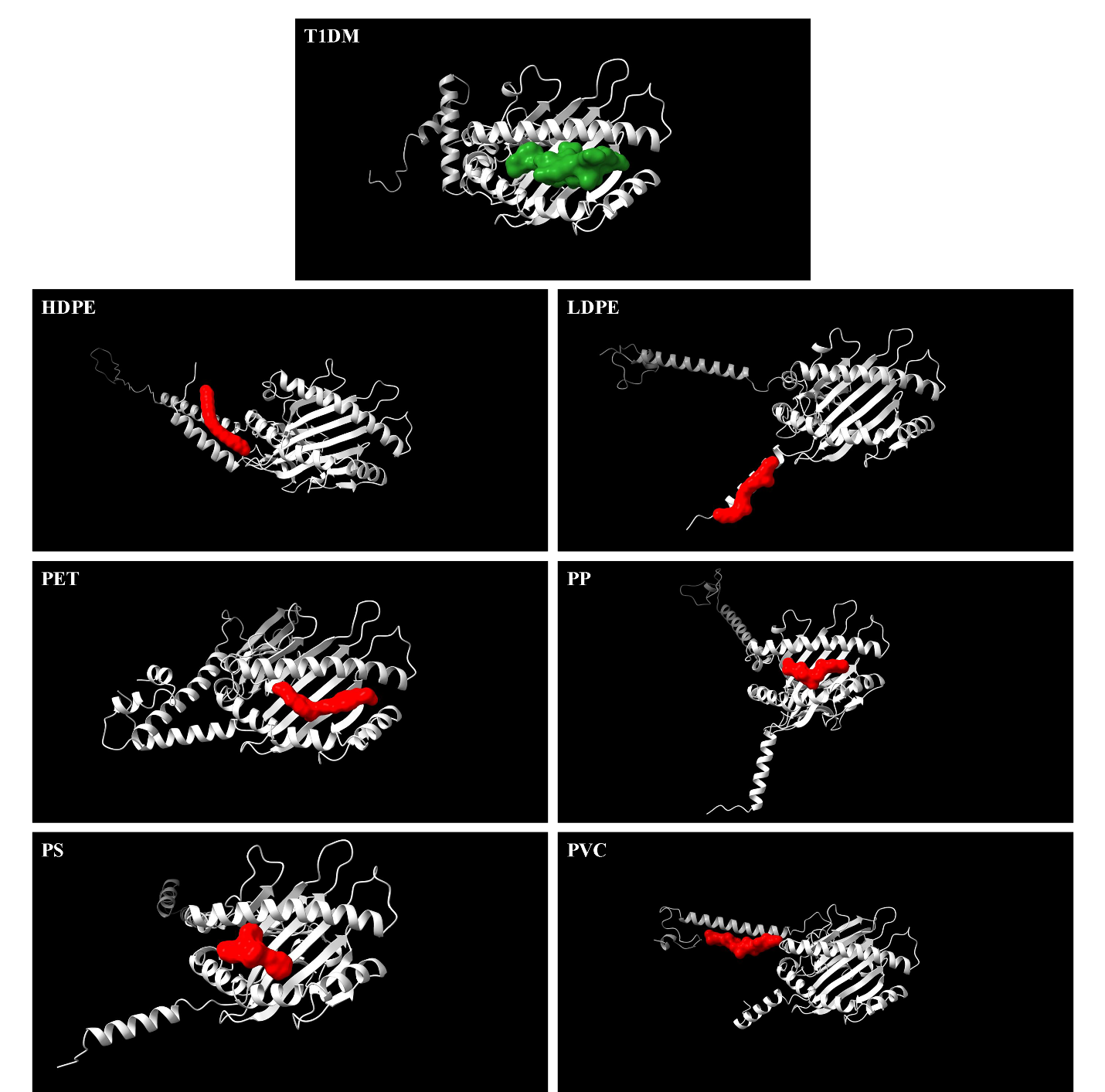

**Figure S2C**: Model structures of all HLA-DQA1*03:01 systems studied here. The T1DM antigen epitopes are depicted in green, and the plastic fragments are depicted in red.

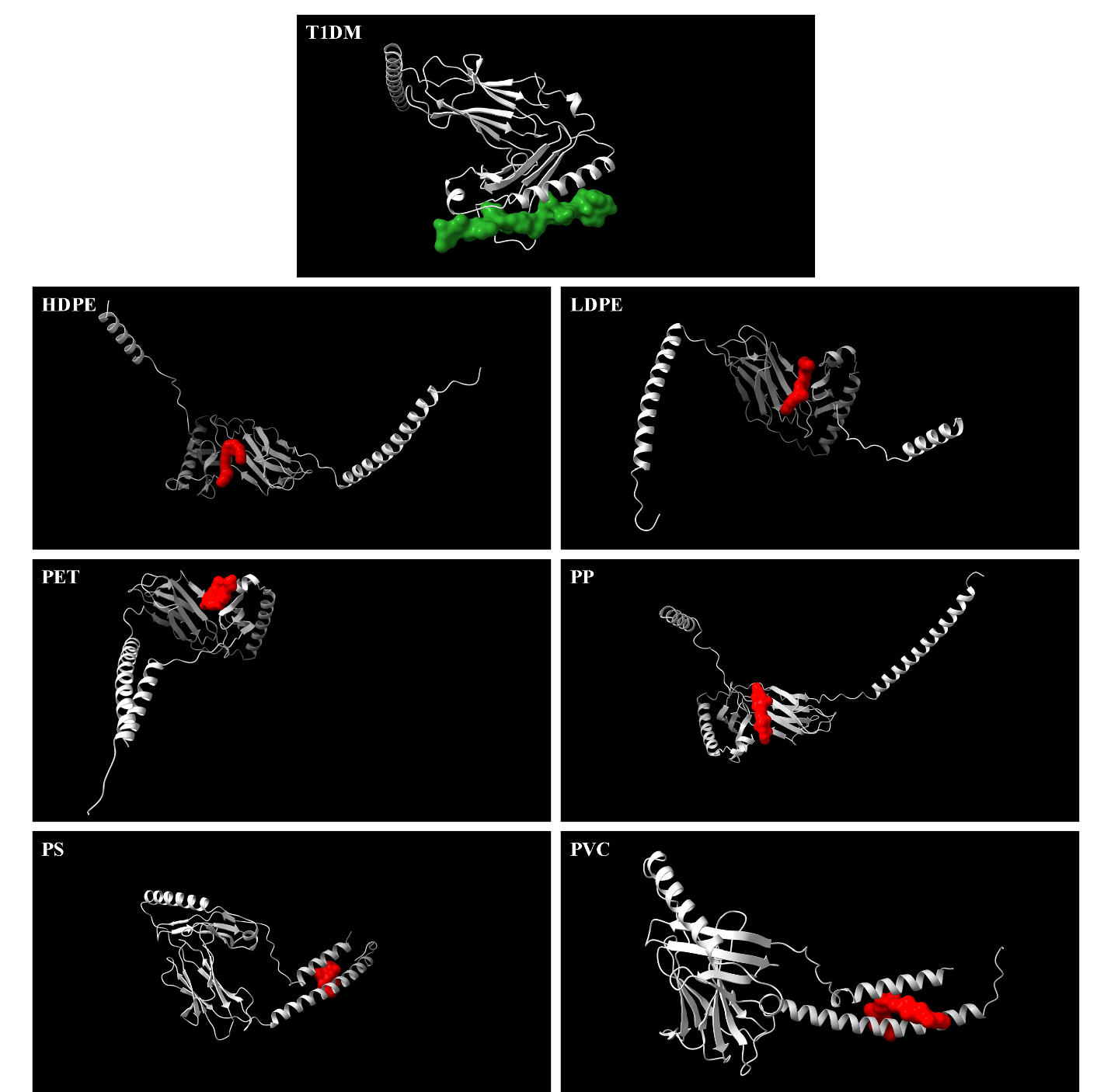

**Figure S2D**: Model structures of all HLA-DQA1*05:01 systems studied here. The T1DM antigen epitopes are depicted in green, and the plastic fragments are depicted in red.

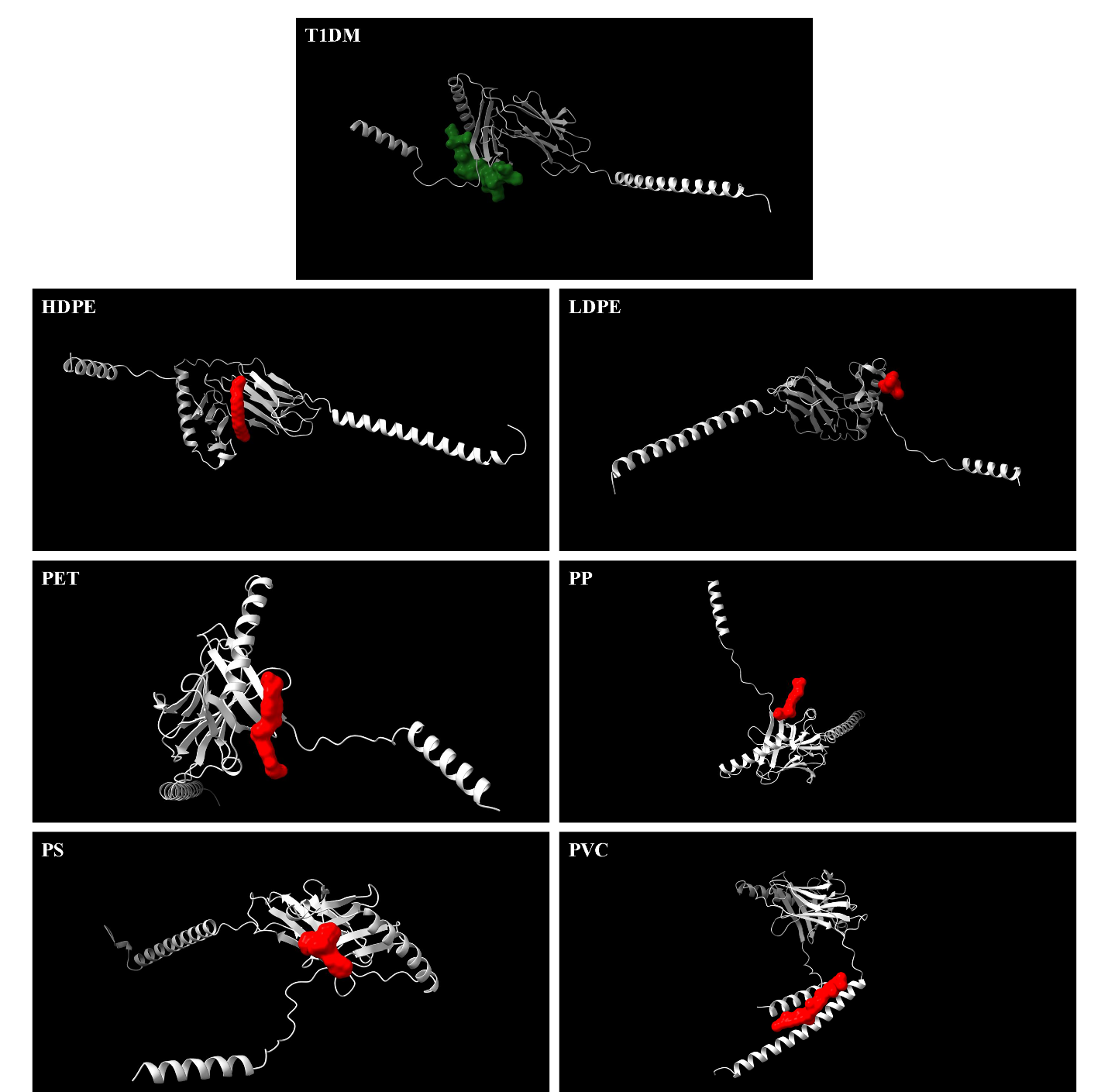

**Figure S2E**: Model structures of all HLA-DQB1*02:01 systems studied here. The T1DM antigen epitopes are depicted in green, and the plastic fragments are depicted in red.

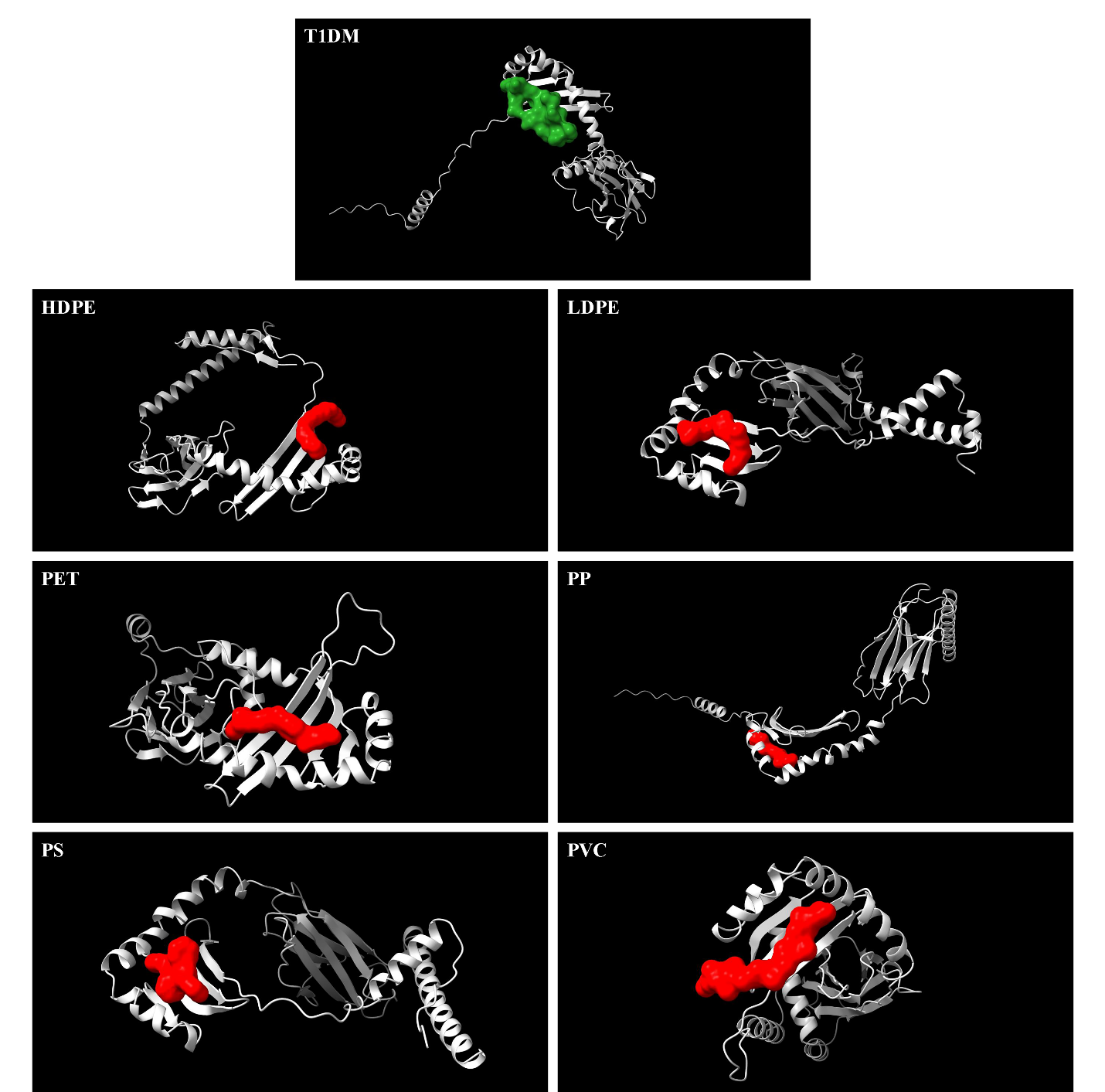

**Figure S2F**: Model structures of all HLA-DQB1*03:02 systems studied here. The T1DM antigen epitopes are depicted in green, and the plastic fragments are depicted in red.

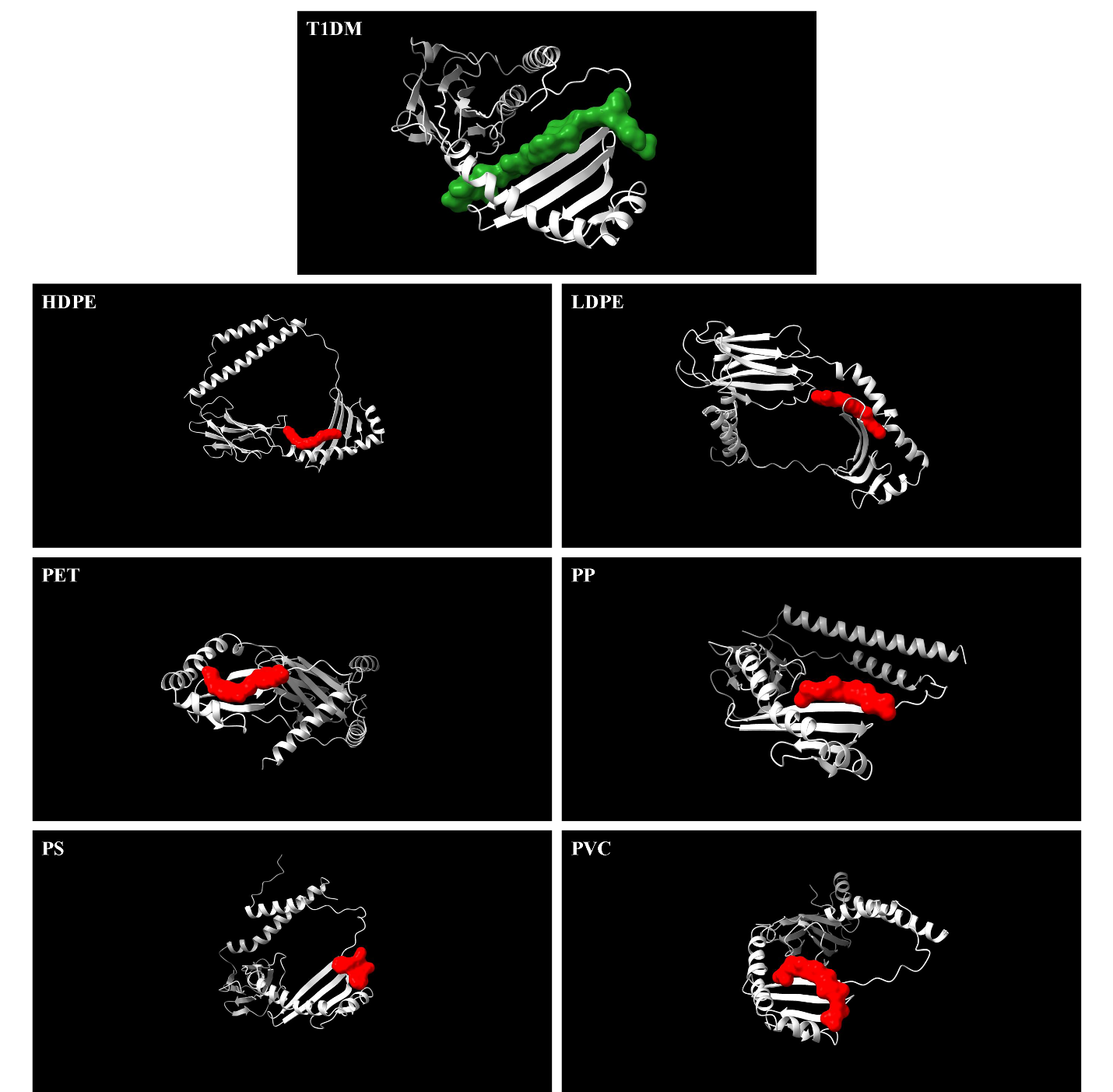

**Figure S2G**: Model structures of all HLA-DRB1*03:01 systems studied here. The T1DM antigen epitopes are depicted in green, and the plastic fragments are depicted in red.

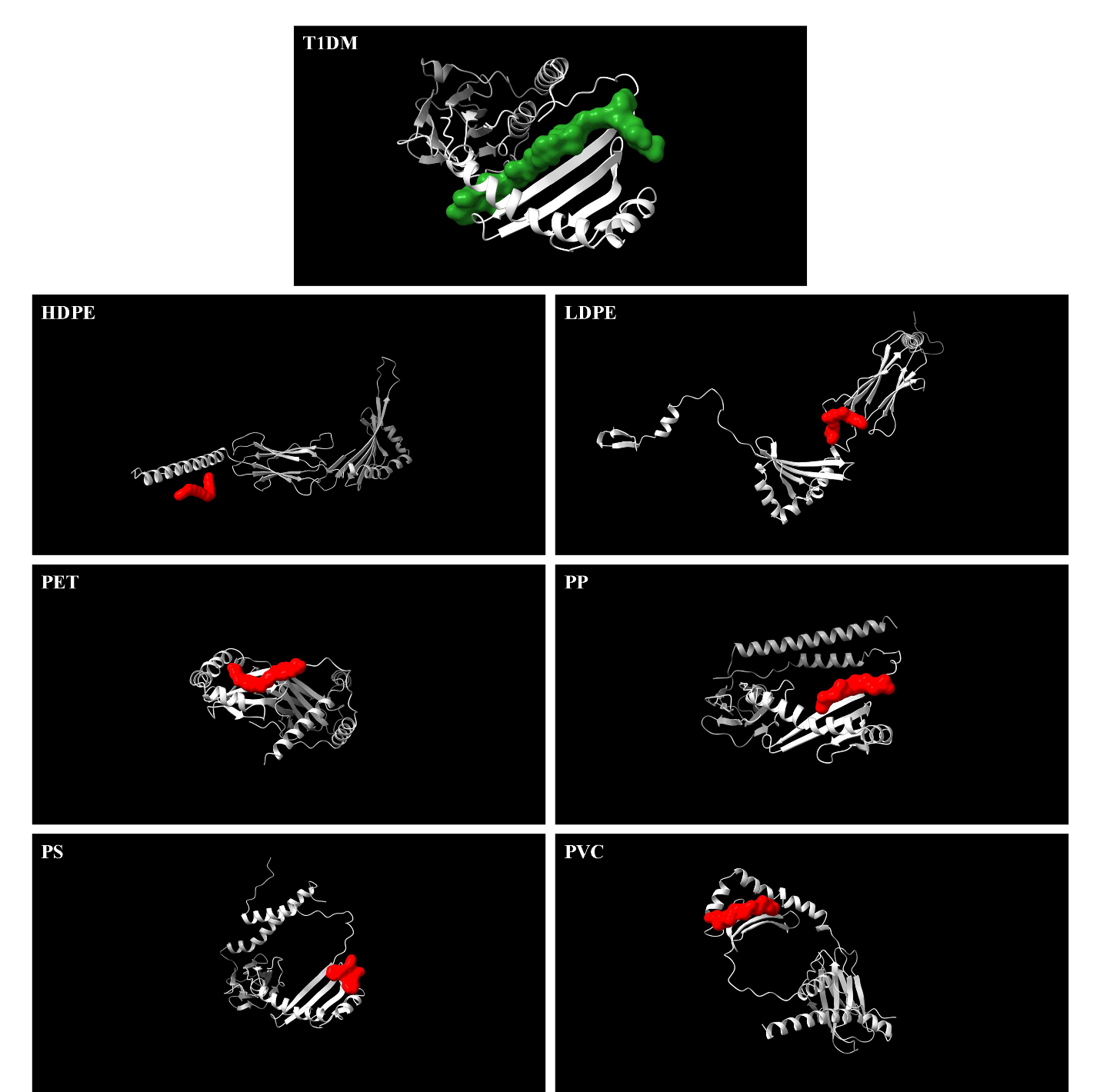

**Figure S2H**: Model structures of all HLA-DRB1*04:01 systems studied here. The T1DM antigen epitopes are depicted in green, and the plastic fragments are depicted in red.

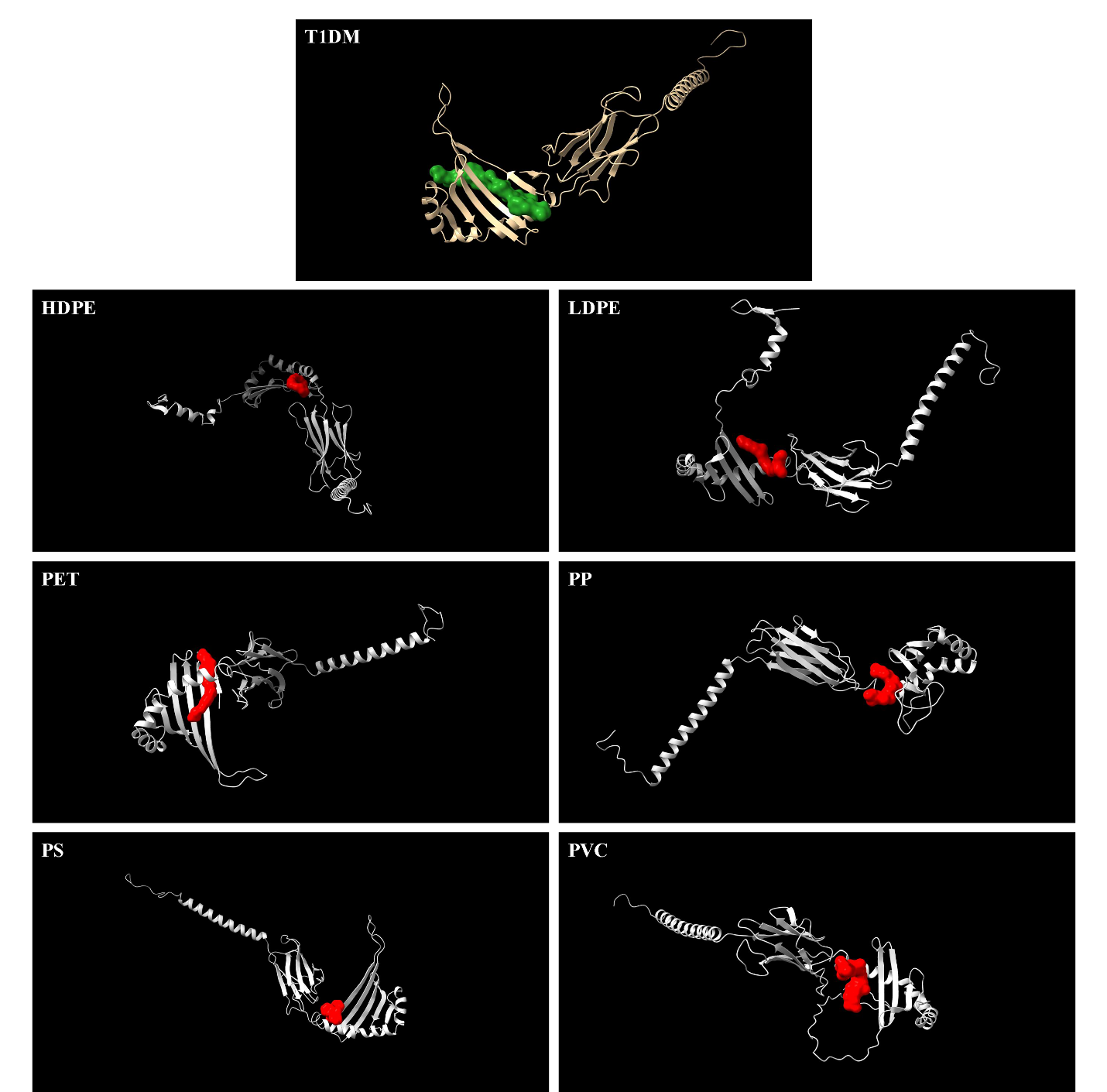

**Figure S2I**: Model structures of all HLA-DRB3*02:02 systems studied here. The T1DM antigen epitopes are depicted in green, and the plastic fragments are depicted in red.

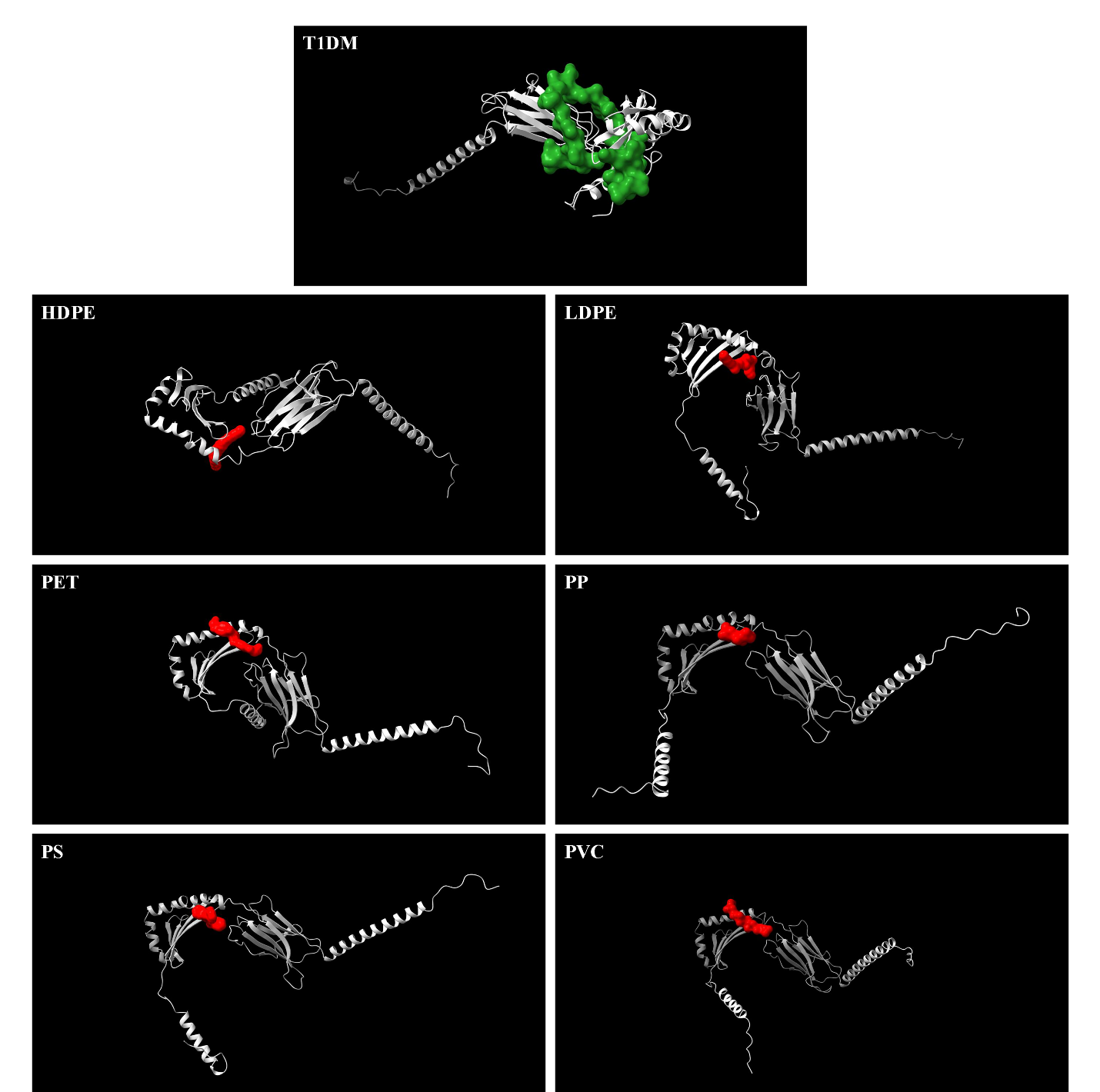

**Figure S2J**: Model structures of all HLA-DRB4*01:01 systems studied here. The T1DM antigen epitopes are depicted in green, and the plastic fragments are depicted in red.

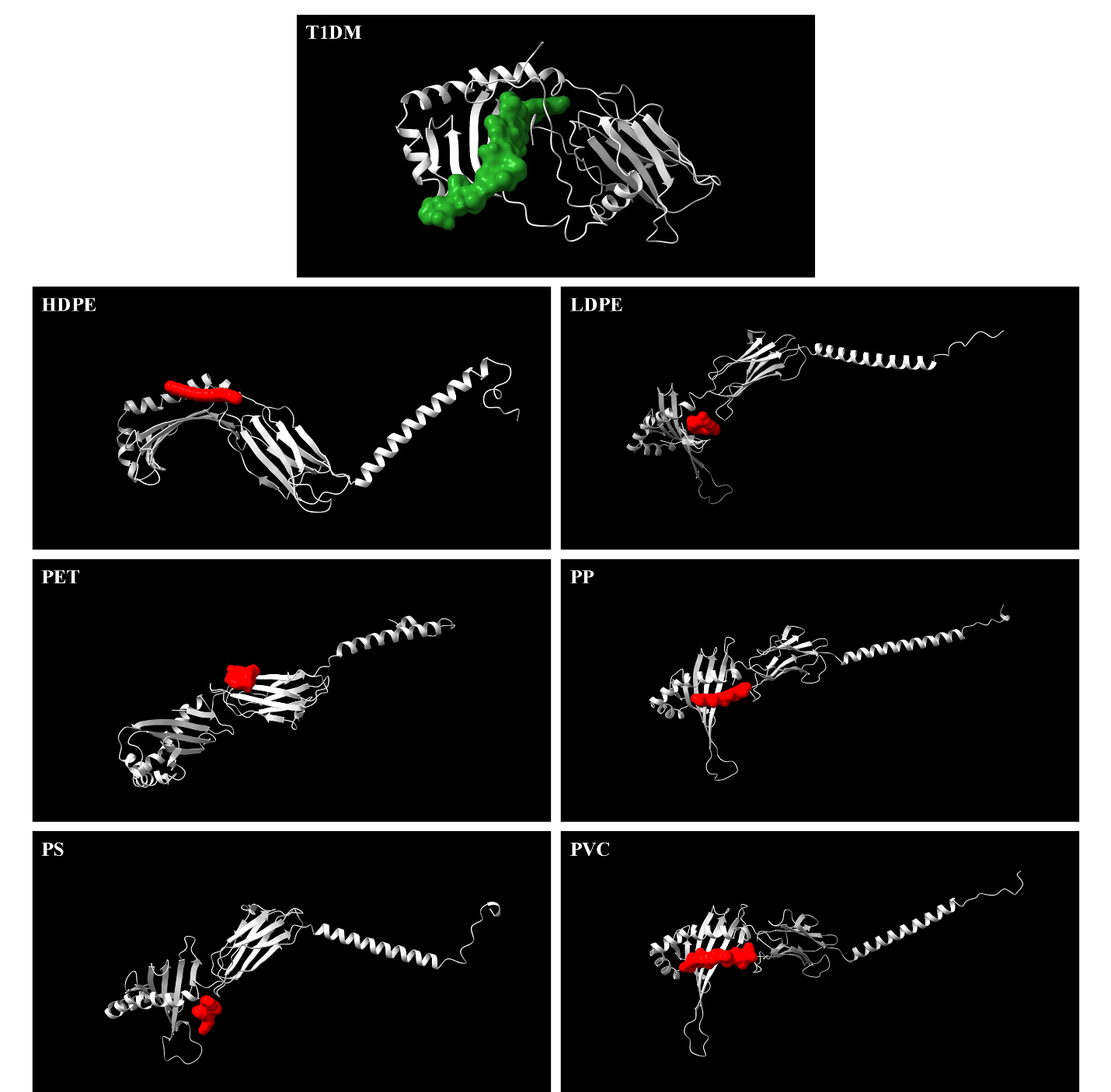
